# Goal-centered representations in the human hippocampus

**DOI:** 10.1101/2021.08.18.456881

**Authors:** Jordan Crivelli-Decker, Alex Clarke, Seongmin A. Park, Derek J. Huffman, Erie Boorman, Charan Ranganath

## Abstract

Recent work in cognitive and systems neuroscience has suggested that the hippocampus might support planning, imagination, and navigation by forming “cognitive maps” that capture the structure of physical spaces, tasks, and situations. Critically, navigation involves planning within a context and disambiguating similar contexts to reach a goal. We examined hippocampal activity patterns in humans during a goal-directed navigation task to examine how contextual and goal information are incorporated in the construction and execution of navigational plans. Results demonstrate that, during planning, the hippocampus carries a context-specific representation of a future goal. Importantly, this effect could not be explained by stimulus or spatial information alone. During navigation, we observed reinstatement of activity patterns in the hippocampus ahead of participants’ required actions, which was strongest for behaviorally relevant points in the sequence. These results suggest that, rather than simply representing overlapping associations, hippocampal activity patterns are powerfully shaped by context and goals.

## Introduction

Every day, people need to plan and execute actions in order to get what they want. Spatial navigation, for instance, requires one to pull up a mental representation of the relationships between different places—i.e., a “cognitive map” (Tolman 1948)—and generate a plan for how to reach a goal. Critically, we can use a cognitive map flexibly, so that, in theory, the same underlying representation can be used to reach different goals. For example, if we wanted to navigate to the Tiger exhibit at the San Diego Zoo we might use the same map-like representation to find the Zebra exhibit.

Several lines of evidence suggest that the hippocampus plays a key role in navigation, though its role in navigation is fundamentally unclear. For example, evidence shows that, during random foraging, “place cells” in the hippocampus encode specific locations within a spatial context (O’Keefe and Dotrovsky, 1971; O’Keefe and Nadel, 1978). However, more recent evidence suggests that the hippocampus is not only important for mapping physical space. Numerous studies have shown that hippocampal activity tracks variables beyond spatial information (Tavares et al., 2015; Park et al., 2019; Aronov et al., 2017). These findings have led to a reimagination of a purely spatial “cognitive map”, whereby the hippocampus can map all manner of spaces (Eichenbaum & Cohen, 2014) and that the hippocampus uses spatial codes in order to encode behaviorally relevant variables (Behrens et al 2018, Stachenfeld et al., 2017, Kaplan, Schuck, & Doeller 2017).

Building on this idea, some have proposed that the hippocampus not only indicates one’s *current* location relative to a map of a physical or an abstract space, but that it also represents possible states or locations that could be encountered in the future (e.g. Mehta et al., 2001, Stachenfeld et al., 2017). According to some versions of the “predictive map” model, through a reinforcement learning process, the hippocampus represents each state in terms of its possible transitions to future states. As a result, the hippocampus enables animals to learn long-term state relationships that would be useful for temporally extended behavior like navigation.

Although there is a substantial body of work on the representation of abstract spaces in the human hippocampus, this work has not completely paralleled work on hippocampal maps in rodents. One key issue identified in single-unit recording studies is that spatial selectivity of place cells is context-specific—that is, the spatial selectivity of a given cell in one environment varies when an animal is moved to a different, but topographically similar environment (O’Keefe and Dotrovsky, 1971, Skaggs and McNaughton, 1998; Leutgeb et al., 2004, Alme et al., 2014, McKenzie et al., 2014). Just as one might pull up different cognitive maps for different physical contexts, it is reasonable to think that we might utilize context-specific maps of abstract state spaces. Theoretical learning models have been proposed to explain how the hippocampus might recognize contexts (Honi et al., 2020, Whittington et al., 2020, George et al., 2021), but there is little empirical evidence showing how context is utilized in abstract spaces.

In addition to context, there is limited information about how goals affect representations of either locations or abstract task states. Most studies of hippocampal place cells in rodents examine activity during random movements through an environment (e.g. Alme et al., 2014), and studies of abstract spaces in humans typically investigate incidental learning of stimulus dimensions or arbitrary state dynamics (Garvert et al., 2017, Schapiro et al 2016, Schuck & Niv, 2016). Research in rodents suggests that representations of space during goal-directed navigation may differ from random or incidental behavior. For example, hippocampal place cells have differential firing fields during planning depending on the future goal of the animal (Ainge et al., 2007; Wood et al., 2000; Ferbinteanu and Shapiro, 2003, Ito et al., 2015), and goal locations tend to be overrepresented by a clustering of spatial codes around them (Dupret et al., 2010, Gauthier et al., 2018). In humans, hippocampal activity patterns during route planning carry information about prospective goal locations in a virtual space (Brown et al., 2016), anticipated stimuli (Garvert et al., 2017), and the magnitude of activity in the hippocampus is modulated by a participant’s distance from a goal location (Patai et al., 2019, Howard et al., 2014). These findings suggest that goals may exert a powerful influence on hippocampal representations.

In the present study, we used functional magnetic resonance imaging (fMRI) to investigate how contexts and goals shape hippocampal representations during planning and navigation. We devised a novel task to investigate the contribution of the human hippocampus to the flexible representation of context dependent goals and plans in an abstract space (Fig. 1). This design allows us to examine the impact of context and goals on hippocampal activity patterns across perceptually similar planned sequences. Critically, we compared evoked patterns elicited for planned sequences that shared a goal to those that had different goals to disentangle the unique contribution of goal information on hippocampal activity patterns. Finally, we analyzed the time course of hippocampal patterns while participants actively navigated during the task to examine if current and future states were reactivated in a way that is consistent with computational models of hippocampal function.

**Fig. 1.**
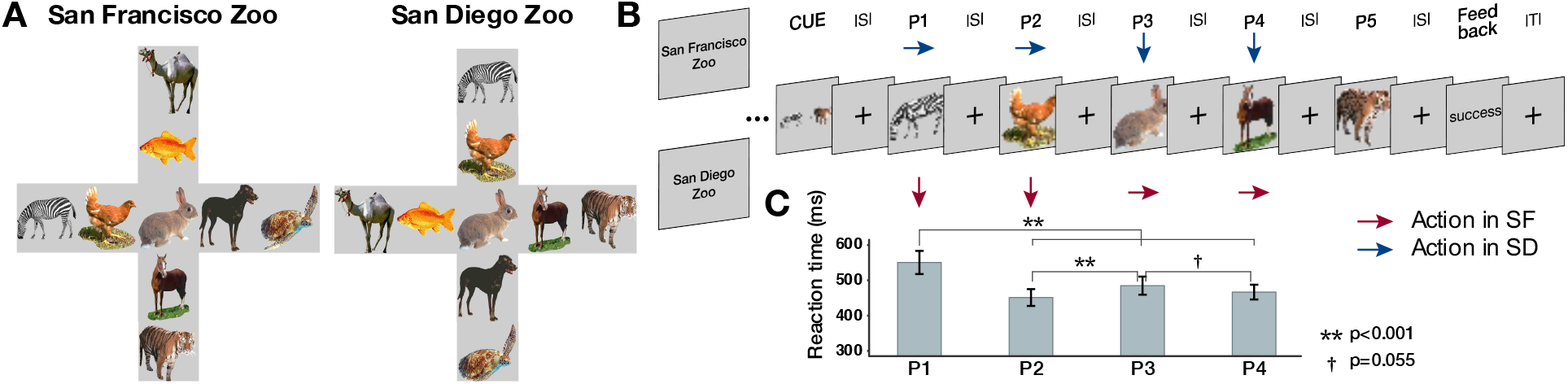
Task Design and Behavioral Results. A) Overhead view of virtual environments. Each context had the same visual information but the specific spatial orientation was mirror reversed and then rotated counter clockwise 90 degrees. This manipulation meant that the action sequence to reach a goal was different across contexts but participants viewed the save visual stimuli. B) Example navigation trial in the scanner. Participants were first cued with a start and goal location and moved navigated through the maze one animal at a time. Inter-stimulus interval (ISI) was 3s. Arrows in red and blue indicate that participants had to make different actions to the same stimuli across contexts to reach their goal during navigation. C) Group level behavioral results from scanner showing elevated reaction times at decision points (Position 1 and Position 3). Error bars represent +-SEM.

## Results

### Navigating an abstract spatiotemporal map

Prior to scanning, participants were trained to criterion (85% accuracy) to navigate to four goal animals in two distinct contexts that consisted of animals that were systematically linked in a deterministic sequence structure (see **Methods**). Each zoo context consisted of the same nine animals arranged in a “plus maze” topology, but the relationships between animals across the two zoos were mirror-reversed and then rotated counterclockwise by 90 degrees (Fig. 1a). At each animal, subjects were able to make one of four button presses that allowed them to transition between animals. In the scanner, participants were asked to use their knowledge of the zoo contexts to actively navigate from a “start animal” to a “goal animal” (Fig. 1b), where start and goal animals were always at the ends of the maze arms. Each trial consisted of a planning phase and a navigation phase. During the planning phase, a cue indicated the start and goal animals. Next, during the navigation phase, participants saw the start animal alone before moving through a sequence of animals to reach the goal animal. For each animal, participants had to decide which direction in the plus maze to move to ultimately reach the goal animal. On any given trial, participants were only allowed four moves to navigate to the goal animal and the interstimulus interval was fixed to ensure that an equal amount of time was spent at each state. In each zoo context, participants planned and navigated 12 distinct sequences (each repeated 4 times across 6 runs of scanning).

Participants were highly accurate at navigating to the goal animal in each context (Context 1: Mean = 93.7%, SD = 12.9%, Context 2: Mean = 94.7%, SD = 12.2%), with no significant differences in accuracy between contexts (t_22_ = 1.16, p = 0.26). This suggests that participants had successfully formed distinct representations of each zoo context. We next tested whether participants’ reaction times would be modulated by differences in the decision-making demands at different locations in the virtual maze. Specifically, our task was structured such that participants were required to initiate their navigation plan at the onset of the start animal (i.e., position one), and at position three – the center of the plus maze, they needed to choose the correct move in order to reach the goal. Accordingly, we expected reaction times (RTs) to be higher at these positions in the navigational sequence than at other positions. Consistent with this prediction, analyses with a linear mixed effects model revealed a significant effect of position (χ^2^(3) = 220.99, p < 0.0001), such that RTs were elevated at position one and position three, relative to other positions (p1 > p2: z = 13.97, p < 0.0001; p1 > p3: z = 9.13, p < 0.0001; p1 > p4, z = 11.67 p < 0.0001; p3 > p2, z = 4.84, p < 0.0001; p3 > p4, z = 2.536, p = 0.0112) (Fig. 1). This shows that decision-making demands at key locations, such as choice points, influenced participants’ response time.

### Hippocampus is sensitive to context-specific sequences in abstract spaces

During the planning phase (i.e., when participants were viewing the cues), we expected that participants should retrieve information about the sequence of state-action pairs that led from the start animal to the goal animal. Our first analyses targeted the extent to which hippocampal activity patterns carried information about the context and the planned sequence. To address this question, we extracted hippocampal multi-voxel activity patterns on each cue trial and calculated pattern similarity (Pearson’s r) between trial pairs that came from repetitions of the same sequence cue in the same context, and compared those to both trial pairs for sequence cues with different start or end points, and trial pairs for sequence cues that came from the same or different context (Fig. 2a). Importantly, visual information was shared across contexts as the cue only indicated the start and goal animal, not the context, and the same cue was associated with different moves between contexts.

**Fig. 2.**
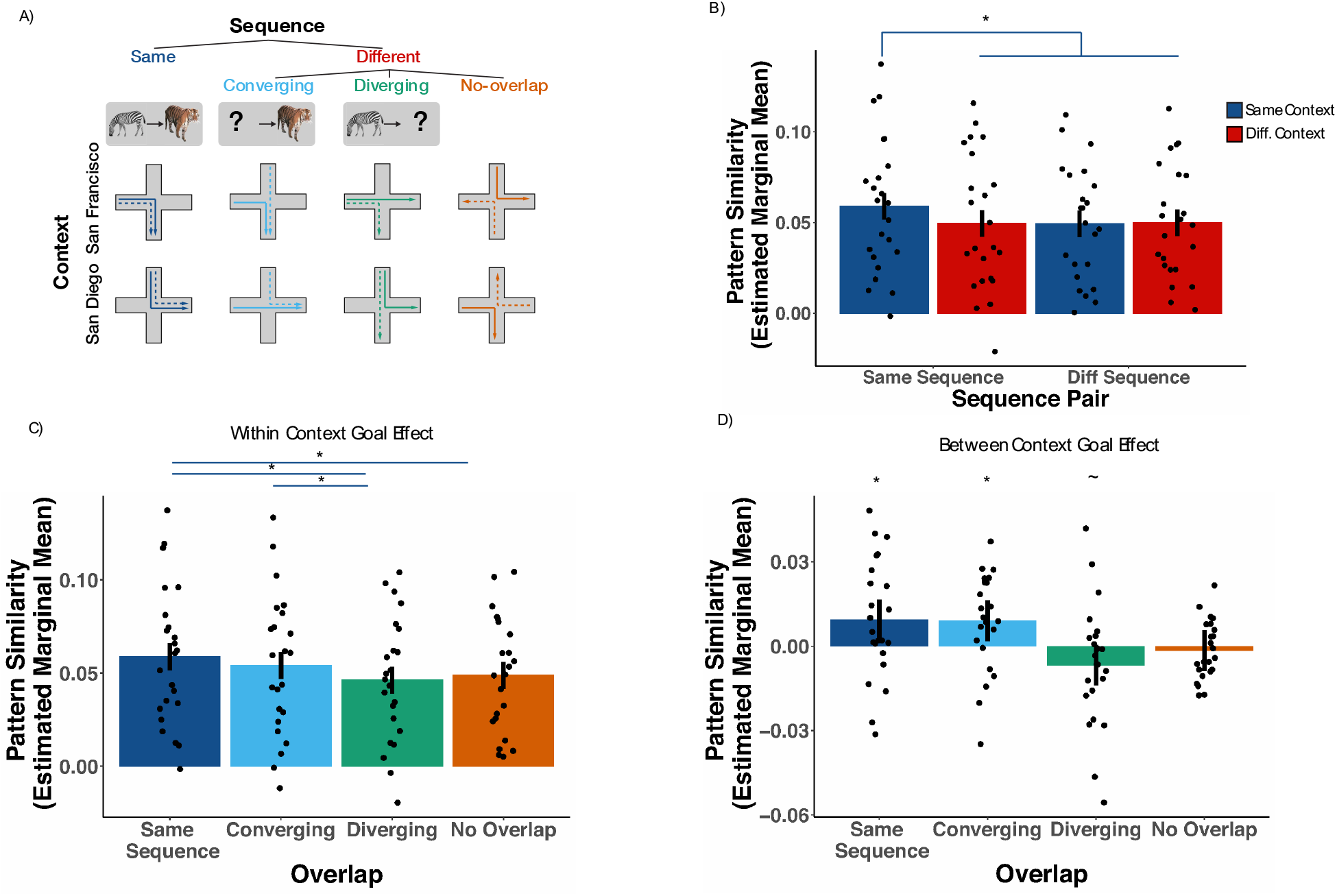
Differential representation of future states in the hippocampus. A) Examples of trial pairs used in pattern similarity analyses during the planning phase. Dashed and solid lines of the same color represent two separate repetitions of the same trial type. B) Results from bilateral hippocampus. Pattern similarity was calculated using estimated marginal means obtained from linear mixed effects models. Pairs of trials sharing sequence and context have significantly higher pattern similarity than all other conditions. C) Pattern similarity results comparing converging and diverging sequences within context. Fully overlapping and converging sequences show higher similarity than diverging sequences. D) Between context overlap effect. Converging and fully overlapping sequences show higher pattern similarity in the same context. Diverging sequences show higher pattern similarity in different contexts. Error bars are 95% confidence intervals of the calculated estimated marginal means. * p < 0.05, ∼p< 0.10.

To test whether hippocampal activity patterns carried information about the context and the planned sequence, we used a linear mixed effects model (Dimsdale-Zucker & Ranganath 2018) with fixed effects of context (same/different) and sequence (same/different), and a random intercept for subject (see Methods for model details and EQ2) to predict pattern similarity in the hippocampus. We reasoned that during planning, participants were retrieving information about the sequence of states and actions to reach their goal. Therefore, we predicted that pattern similarity should be higher for sequences that shared the same state-action pairs. Moreover, we predicted that this effect should be context-specific, as the same sequence across contexts have different state-action pairs. Consistent with this prediction, we found a significant sequence by context interaction (Fig. 2b: χ^2^(1, N = 23) = 4.26, p = 0.04). Follow up tests showed that patterns evoked by the same sequence cue in the same context were significantly different than all other trial pairs (same seq. + same cx. > diff. seq. + same cx.: z = 2.77, p = 0.006; same seq. + same cx. > same seq. + diff. cx.: z = 2.73, p = 0.006; same seq. + same cx. > diff. seq. + diff. cx.: z = 2.61, p = 0.009; see Fig. 2b). These results show that hippocampal activity patterns carried information about planned state-action sequences within specific contexts.

### Hippocampal activity patterns reflect future goals during planning

The above analysis demonstrates that hippocampal activity patterns carry context-specific information about planned sequences, consistent with prior work showing that the hippocampus represents information about specific sequences of objects (Hsieh et al., 2014; Schapiro et al. 2016; Kalm, Davis, & Norris, 2013; Agster, Fortin & Eichenbaum 2002), as well as work suggesting that the hippocampus forms distinct representations of overlapping spatial routes (Chanales et al., 2017, Wood et al. 2000). However, there are reasons to think that hippocampal sequence representations might become more similar under certain circumstances. For instance, if the hippocampus uses predictive maps that carry information about possible future states (Stachenfeld et al., 2017), one might expect similar representations of sequences that share the same starting point but lead to different goals by more heavily weighting the immediate state-action pairs that follow planning (“diverging sequences”; see Methods for successor representation simulation details and **Fig. S1**). On the other hand, it is possible that goals are more heavily weighted during planning (Mattar and Daw 2018), and thus we might expect similar representations of sequences that lead to the same goal but start at different states (converging). We sought to test these ideas by comparing pattern similarity during cues associated with repetitions of the same sequence, cues associated with different sequences that converged on the same goal (converging), cues associated with sequences that shared the same starting state but with different goals (diverging), and cues associated with non-overlapping sequences (no overlap)(Fig. 2a).

A linear mixed effects model with fixed effects for overlap (same sequence/converging/diverging/no overlap) and context (same/different) and a random intercept for subject (see Methods for model details and EQ3) showed a significant context by overlap interaction (χ^2^(3, N = 23) = 14.75, p = 0.002). (Fig. 2c and 2d). Follow up tests revealed that, within a context, cues with converging goals had significantly higher pattern similarity than cues with diverging goals (z = 2.19, p = 0.03), and same sequence cues had higher pattern similarity than cues with diverging goals (z = 3.49, p = 0.0005) and sequence cues with no overlap (z = 2.77, p = 0.0056). However, converging sequences were not significantly different from the same sequence (z = 1.30, p = 0.194). In sum, these results show that during planning, representations in the hippocampus are differentiated based on future context-specific goals, suggesting that goals fundamentally shape representations in hippocampus via shared patterns between sequences that lead to the same goal.

### Differences in pattern information during the cue period cannot be explained by shared motor plans or sensory details

The present results are consistent with the idea that the hippocampus supports planning of state-action sequences toward a goal. Importantly, our cues were carefully controlled, such that subjects were viewing visually identical stimuli across contexts and did not make responses during the planning phase. However, some factors could have covaried with our manipulations. One possible explanation is that, both within and across contexts, sequences can involve the same motor responses. While subjects are not actively making movements during planning, the pattern of results in hippocampus could be driven by overlap in motor planning in converging vs. diverging sequences. To ensure context effects observed in hippocampus were not due to shared motor information during planning, we examined trial pairs that had the exact same moves, trial pairs that had two moves in common, and pairs that had no moves in common to ensure that movement information alone was not modulated by context in the hippocampus. Results showed no effect of planned moves or context on pattern similarity (main effect of context: χ^2^(1, N = 23) = 0.46, p = 0.5; main effect of move: χ^2^(2, N = 23) = 1.56, p = 0.46; interaction: χ^2^(2, N = 23) = 2.68, p = 0.26) **Fig S2**.

As a positive control analysis, we also examined an anatomically defined motor cortex ROI (BA4a/4p) to investigate whether we could detect sensorimotor representations and if they were modulated by context information during planning. Results revealed a significant main effect of planned move (χ^2^(2, N = 23) = 40.40, p < 0.0001), and importantly showed that planned movement was not modulated by context (main effect: χ^2^(1, N = 23), = 0.01 p = 0.94; Interaction: χ^2^(2, N = 23), = 1.26, p = 0.53 **Fig S2**)(See Methods and EQ4 for model details). Taken together, these results show that our cue period results in the hippocampus cannot be solely explained by shared motor information of a plan and highlights the role of the hippocampus in retrieving the specific state-action sequence required to execute a given plan.

To mitigate against a potential confound that low-level visual features could explain the observed effects of context in our hippocampal results (Huffman and Stark, 2017), we ran a control analysis on an anatomically defined visual cortex ROI (V1/V2). To do this, we compared pattern similarity between trials where the cue image had the same images, one image in common, or no images in common. This analysis is identical to the overlap analysis run on hippocampus above (see Methods and EQ3 for model details). We found that this visual cortex ROI was only sensitive to visual information (Main effect of overlap – χ^2^(3, N = 23) = 90.24, p < 0.0001 and not context (χ^2^(1 N = 23) = 0.05, Interaction: p = 0.82; χ^2^(3, N = 23) = 0.76, p = 0.86 **Fig. S2**). These results suggest that our main HC findings represent something beyond a shared sensory representation. Altogether, these analyses provide an important control and bolster our interpretation of the findings from our analyses of the hippocampus, by showing that primary sensory areas are activating behaviorally-relevant representations during planning, but that the effects of context and goal are only present in hippocampus.

### Representation of behaviorally relevant sequence positions during navigation

Having established that the hippocampus represents information about context-specific goals during planning, our next analyses turned to how state-action information is dynamically represented during navigation. To examine if hippocampal activity patterns carried information about current or future states during navigation, we extracted the time-series for each navigation sequence using a variant of single trial modeling that utilizes finite impulse response (FIR) functions (Turner et al., 2012), allowing us to examine activity patterns for each time point (TR) as participants navigated through the sequence of items. As depicted in Figure 3, we quantified pattern similarity between pairs of navigation sequences (e.g. zebra to tiger sequence compared to camel to tiger sequence) at different timepoints (e.g., TR 1 to TR 10), which yielded a timepoint-by-timepoint similarity matrix for each condition (converging or diverging sequences). The diagonal elements for this matrix reflect similarity between pairs of animal items from the same timepoint in the sequence. Off-diagonal elements reflect the similarity between an animal at one timepoint in the sequence and animal items at other timepoints in the sequence. This technique is conceptually similar to cross-temporal generalization techniques used in pattern classification analyses (King & Dehaene, 2014).

**Fig 3.**
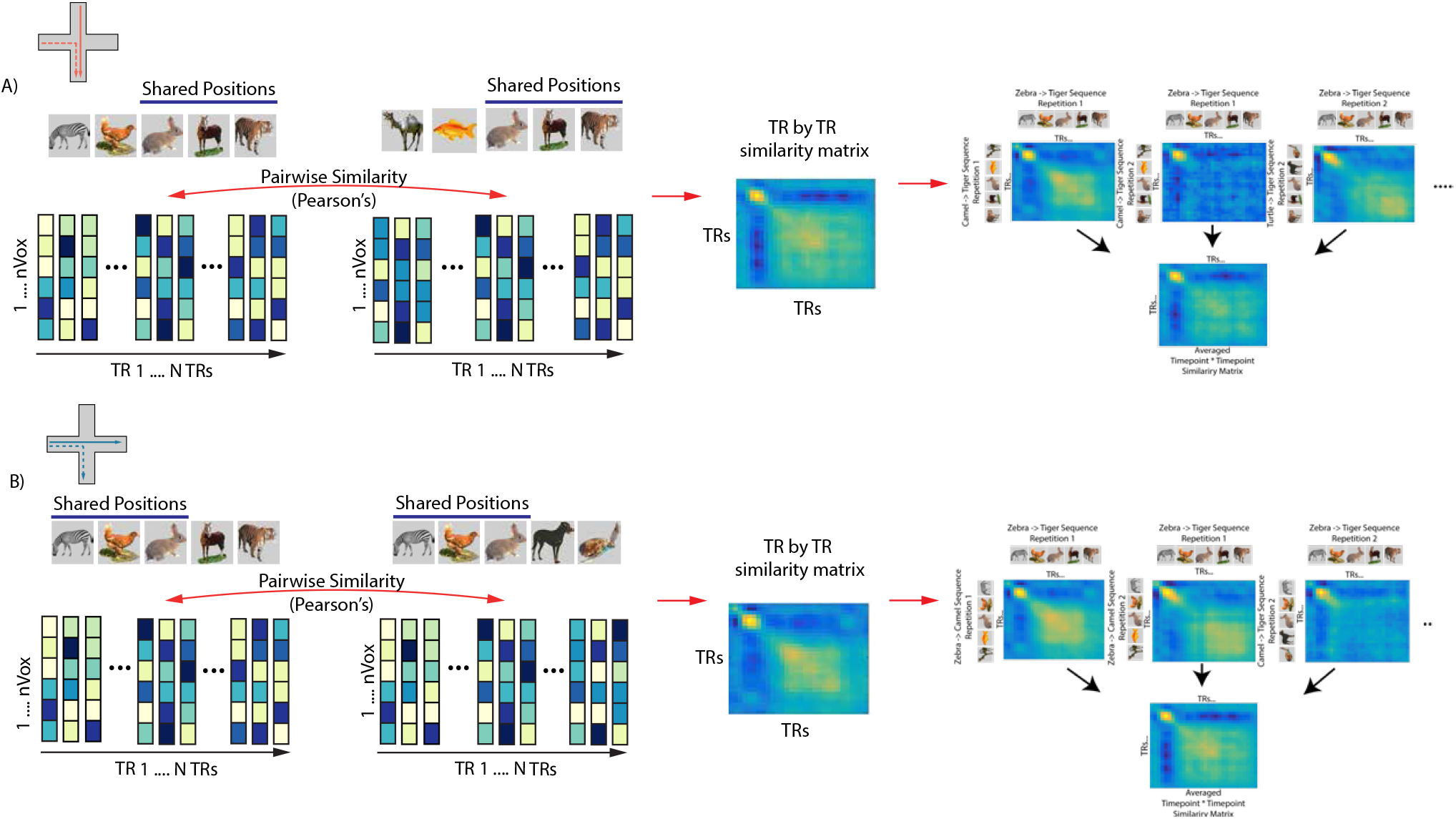
Schematic depiction of procedure to obtain time point by time point similarity matrices. A) (Left) Dashed and solid lines on the maze indicate an example pair of trials correlated. TR by TR spatio-temporal patterns were obtained for a pair of sequences (converging in this example). Pattern similarity was computed between every possible pair of spatial patterns (voxels) over all timepoints (TRs) from a region of interest. (Middle) This procedure yielded a TR by TR similarity matrix for a given sequence pair. Note that because the sequences are from different repetitions across fMRI scanning runs the diagonal is not perfectly correlated. (Right) This was repeated for every possible converging sequence pair in the data set. The resultant TR by TR matrices were than averaged to create a subject level converging TR by TR matrix. Subject-specific averaged TR by TR matrices were than statistically compared to diverging sequences using cluster-based permutation tests (see Methods). B) Same as A but using an example diverging sequence pair.

Pattern similarity analyses during navigation specifically focused on converging and diverging sequence trials. Converging and diverging sequences were chosen because these sequences have an equal number of overlapping states, but the timing of the overlap is systematically different. By computing pattern similarity differences between converging and diverging trials, we were able to assess the representation of current and future states in the hippocampus.

One possibility is that hippocampal patterns during navigation should only be sensitive to what you are seeing or doing at a given moment, similar to position or place codes. In this case, we would expect pattern similarity differences between converging and diverging sequences close to the diagonal of the timepoint-by-timepoint matrices — that is, we would expect higher pattern similarity for diverging pairs during timepoints early in the sequence and higher pattern similarity for converging pairs during timepoints late in the sequence.

Alternatively, other studies have shown that the hippocampus represents not only the current location but also temporally or statistically related states (Stachenfeld et al., 2017, Garvert et al., 2017, Schapiro et al., 2016). In this case, at early positions, we would expect higher pattern similarity for diverging sequences, both on-and off – diagonal, and at late positions, we would expect higher pattern similarity for converging sequences both on-and off –diagonal.

A third possibility is that the hippocampus might preferentially represent goal-relevant information during navigation (Mattar et al., 2018). Consistent with this idea, work in rodents has shown the hippocampus reactivates neural ensembles associated with both future and past events at critical decision points (Diba, Buzaki 2007, Johnson and Redish, 2007, Carr et al., 2011, Pfeiffer and Foster, 2013). In our study, the most behaviorally important points in a navigated sequence were the starting point (position 1), when a goal-directed plan must be initiated, the center of the maze (position 3), a critical sub-goal where one’s decision will determine the ultimate trial outcome. This was confirmed by our behavioral analyses that revealed that participants were slower to respond at positions 1 and 3 (Fig. 1). We therefore reasoned that participants might be likely to prospectively retrieve hippocampal representations of these states during navigation. In this case, we would expect to see higher off-diagonal pattern similarity in converging sequences, such that across converging sequences, activity patterns associated with goal states would be correlated with activity patterns during earlier positions in the sequences. This effect should be higher for converging sequences than for diverging sequences because activations of state-action pairs should become more similar later in the sequence. Conversely, diverging sequence state-action pairs should be more similar earlier in the sequence but then decrease as the sequence progresses.

We calculated timepoint-by-timepoint correlation matrices (Pearson’s r) in hippocampus for converging and diverging sequences then statistically compared them using cluster-based permutation tests (10,000 permutations). We found several clusters showing higher similarity for converging compared to diverging sequences (Fig. 4). Interestingly, there was a significant off-diagonal cluster (outlined in red: p = 0.038, corrected) that roughly corresponded to the first item in the sequence (position 1) activating the central position (position 3) (approx. TRs 10-15). Other clusters tended to overlap with key locations in the experiment, which roughly correspond to position one activating position five (TRs 18 to 21) and position three activating position five (TRs 18 to 20) (Fig. 4e), although these clusters did not survive multiple comparison correction. These data suggest that hippocampus plays a phasic role in the activation of patterns that contain information about future states and prioritizes both goal and sub-goal information during active navigation. These findings, along with the planning period results, suggest that during goal-directed behavior, the representation of certain states is modulated in the hippocampus depending on its importance for the current plan.

**Fig. 4.**
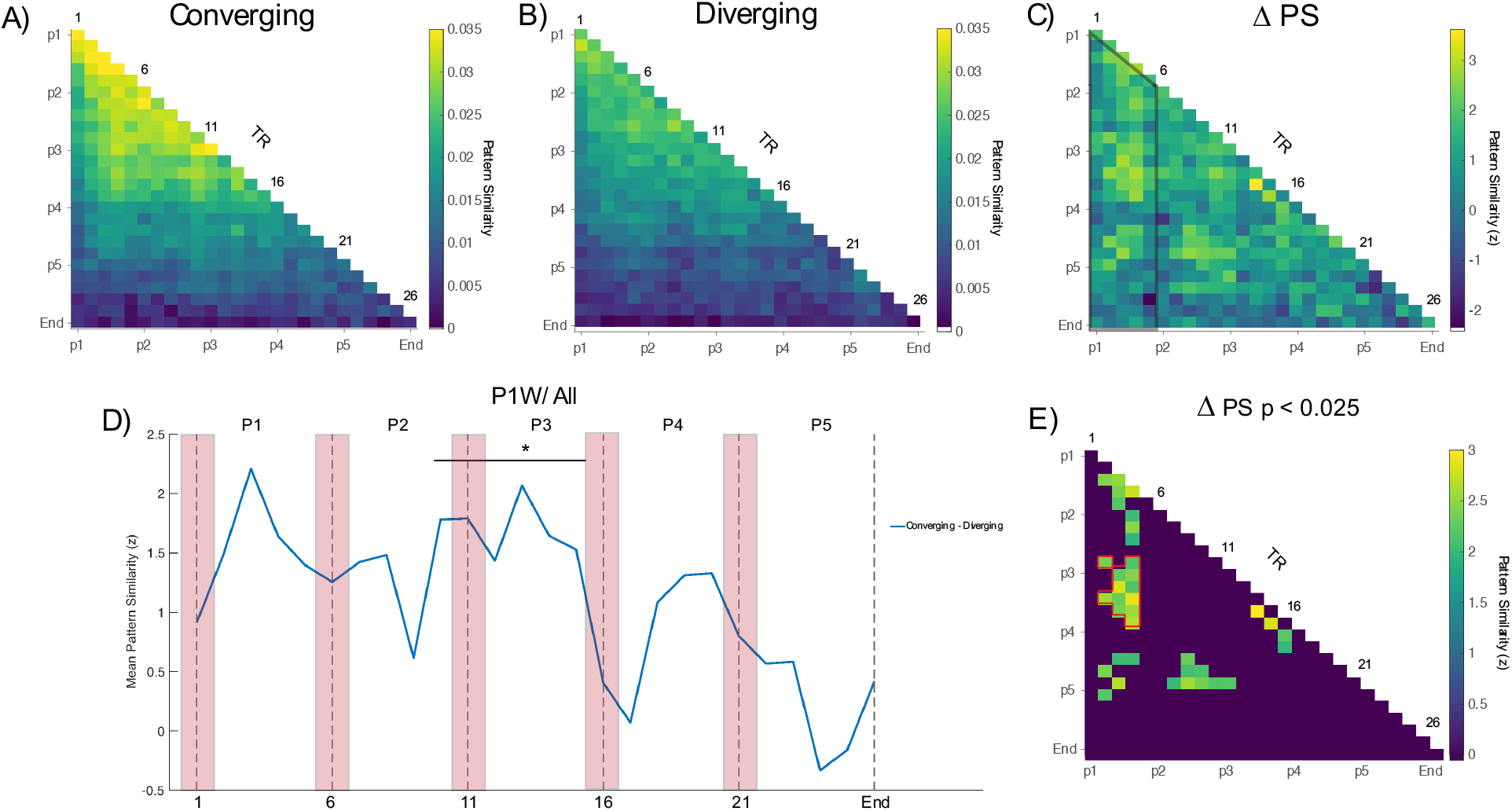
Results from TR by TR pattern similarity analysis during active navigation in bilateral hippocampus. A) Group level pattern similarity results from converging sequences during active navigation. B) Same as A but showing diverging sequences. C) TR by TR pattern similarity results depicting a statistical map of converging – diverging. Z values were calculated using a bootstrap shuffling procedure with 10,000 permutations. Opaque black lines indicate region where data were averaged for visualization purposes. D) Group level time course obtained by averaging over time points outlined in black in C. Horizontal black bar illustrates the approximate time points of the significant cluster in E. E) Thresholded statistical map at p < 0.025. Cluster based permutation tests with 10,000 permutations (Maris and Oostenveld, 2007) were performed with a cluster defining threshold of p < 0.025 and a cluster alpha of 0.05 Outlined in red is a significant cluster of timepoints that survives multiple comparisons correction (cluster mass = 29.44, p = 0.038). Note that this cluster corresponds to approximately position 1 activating position 3 which was shared by both converging and diverging sequences.

## Discussion

The goal of the present study was to identify how the hippocampus represents task information during planning and navigation towards a behavioral goal. During planning, we show that hippocampal representations carried context-specific information about individual sequences to a goal. Surprisingly, not all sequences were equally differentiated, such that sequences that converged on a common goal showed higher pattern similarity compared to diverging sequences, despite an equal amount of overlap between the conditions. These results suggest that the hippocampus forms integrated representations of sequences that lead to the same goal states, and that it differentiates between sequences that start from the same state but lead to different goals. Similarly, during navigation, we found that the hippocampus prospectively activated information about upcoming states and that this effect was strongest in relation to key decision points and goals. Our results are consistent with the idea that, rather than simply representing overlapping associations, hippocampal representations are powerfully shaped by context and goals.

### The hippocampus represents context-specific goal information during planning

A key finding from the present study is that, during planning, hippocampal activity patterns are organized such that they either generalize or differentiate between sequences depending on the goal, and do so in a context-specific manner. These findings are relevant to theories which propose that prospective thought (prediction/planning) relies on the same circuitry used for episodic memory (Hassabis et al., 2007; Schacter et al., 2007, Addis et al., 2012). In support of this idea, sequential place cells activate along the path an animal will take in a phenomenon described as “forward replay” (Johnson and Redish, 2007, Pfieffer and Foster 2013). Building on this work, Brown et al., (2016) used high-resolution fMRI in humans to examine hippocampal activity during goal-directed navigation in a virtual reality (VR) paradigm. Brown et al. demonstrated that, during planning, hippocampal activity patterns could be used to accurately decode future navigation goals, even across different start positions and routes. Thus, their findings demonstrated that fMRI activity patterns in the hippocampus carried information about future navigational goals. Brown et al. (2016) interpreted their findings as evidence that the hippocampus supports imagination or mental simulation of a route towards a goal.

Our findings suggest an important constraint on the role of the hippocampus in imagination and simulation. In our study, if participants simulated the sequence of sensory events that led to the goal (i.e., imagining the sequence of animals), we would expect hippocampal representations to generalize across repetitions of the same sequence of animals across the two different zoo contexts. Instead, we found that hippocampal representations during planning were context specific, such that pairs of trials involving the same sequence of animals across different contexts were indistinguishable from entirely different sequences. Moreover, similarity across different sequences that led to the same goal in the same zoo context was indistinguishable from similarity across trial pairs involving the same sequence in the same context. Thus, in our study, hippocampal activity most likely did not reflect imagination of a sequence of stimuli per se, or even a specific sequence of states, but rather a more abstract representation of the context and goal state.

Together with prior research, our results are relevant to an emerging body of work suggesting that goals and other salient locations exert a powerful force on spatial and non-spatial maps in the brain (McKenzie et al., 2013, 2014; Boccara et al., 2019; Butler & Hardcastle et al., 2019; Brunec et al., 2018). For example, McKenzie et al., (2014) found that rewarded events had higher pattern similarity within a context compared to unrewarded events. Moreover, there is evidence that, after learning in a reward-based foraging task, place cells tend to be clustered around goal locations (Dupret et al., 2010, Gauthier et al., 2018). This could go some way towards explaining our results of increased pattern similarity for sequences that converge on the same goal, while context-specific remapping of place cells is also in line with our results of context-specificity. Considering the current work and past findings, we propose that hippocampal representations are flexibly modulated depending on current behavioral demands, incorporating trial-specific information that allows organisms to realize a specific goal (Ekstrom and Ranganath, 2017).

### Reinstatement of remote timepoints in the hippocampus during navigation

If the hippocampus supports prospective planning for goal-directed navigation, then it is important to understand how it functions when such actions are taken when navigating abstract spaces. For example, if the hippocampus is involved in retrieving the specific state-action plan, what is its function once this plan is executed? To address this question, we contrasted pattern similarity during the navigation phase across pairs of converging sequences against pairs of diverging sequences. By comparing navigation activity patterns from each time point against later time points, we were able to detect pattern similarity between current and remote timepoints that is consistent with a reinstatement like phenomenon (Günseli et al., 2020).

As noted above, the animals in the first three positions overlapped across diverging sequences, whereas the animals in the last three positions overlapped across converging sequences. Thus, if the hippocampus only represented the current state during navigation, we would have expected pattern similarity on the diagonal in Figure 4 to be higher for diverging trials for early time points, and then higher for converging trials in the later time points (see also **Figure S4**). Instead, we found that the significant differences between converging and diverging trial pairs were primarily off of the diagonal, suggesting that, during the navigation phase, hippocampal patterns carried information about behaviorally relevant remote timepoints along the route^1^. This effect was seen across sequences that converged on the same goal, which all require the same decision at position three, although these sequences differed at positions one and two. As such, the relationship between patterns at position one and remote time points is likely driven by the prospective activation of future states likely corresponding to the next key decision point at the center of the plus maze (position three) and the final goal (position five).

Consistent with our study, research in rodents shows that hippocampal ensemble activity differs between routes that share a common path but lead to a different goal (Frank et al., 2000; Wood et al., 2000, Ferbinteanu and Schapiro, 2003; Ito et al., 2015; Markus et al., 1995). In addition, there are several reports of goal locations and future states being represented in human hippocampus (Watrous et al., 2018; Ekstrom et al., 2003; Brown et al., 2016). Research in rodents shows that in navigation and decision– making tasks place cell activity is activated in forward sweeps allowing an animal to simulate the future (Johnson and Redish, 2007; Pfeiffer and Foster et al., 2013). In humans, there are several reports of predictive hippocampal representations that are related to future behavior in both spatial and non-spatial tasks (Momennejad et al., 2018, Garvert et al., 2017, Schapiro et al., 2016, Brown et al., 2016). Our data add to theories of predictive representations in the hippocampus by suggesting that the hippocampus may emphasize strategically important states for reinstatement during ongoing behavior. Thus, we propose that during goal-directed behavior, the human hippocampus does not solely reflect the current state during navigation, or only the immediate future, but predicts states for key decision points along the route – here the center of the maze and the goal. These results align with recent findings in rodents and computational models showing place cells associated with behaviorally relevant locations in an environment are preferentially reactivated (e.g. Mattar and Daw, 2018).

A specific computational implementation of a predictive map model, the successor representation, states that the hippocampus is involved in learning relationships between states and actions, and that its representations reflect expectations about future locations (Stachenfeld et al., 2017; Mommenejad, 2020). We used this computational model to generate simulated pattern similarity results, and surprisingly, these simulated matrices were qualitatively different from what we observed in the hippocampus.

In our simulations, the successor representation reflected the transition probabilities between states, such that adjacent states were more similar than non-adjacent states. This is because, participants transitioned between all start and end positions equally in both directions. Thus, the model could not reproduce the difference between converging and diverging sequences either during the planning or navigation phases. More generally, it is possible that state space representations in the hippocampus will not always reflect one algorithm such as the SR. Instead, hippocampal representations of physical space (Ekstrom and Ranganath, 2017) and abstract state spaces (Boorman, Sweigert, & Park, 2021) are likely to be more flexible, reflecting the computational demands of the planning problem, the subject’s experience with the problem, and the situation. In the present study, the task might have encouraged a model-based planning strategy, in which future goals and key states are strategically retrieved and represented in hippocampus. In other tasks, where the structure is well learned and people do not need to re-plan, hippocampal state spaces might resemble successor-based maps.

## Conclusion

Human behavior is characterized by the ability to plan and flexibly navigate decision spaces in order to realize future goals. The present study demonstrates that the human hippocampus generates context-specific, goal-centered representations that can be flexibly called upon in the service of goal-directed planning and navigation. These findings can contribute to the development of unified models accounting for hippocampal contributions to memory, navigation, and goal-directed sequential decision-making (Eichenbaum, 2017; Wikenheiser & Schoenbaum 2016; Bellmund et al., 2018). Additionally, this work highlights the importance of studying goal-directed behavior, attentional modulation of memory representations, and their consequences on planning.

## Methods

### Participants

Thirty healthy English-speaking individuals participated in the fMRI study. All participants had normal or corrected-to-normal vision. Written informed consent was obtained from each subject before the experiment, and the Institutional Review Board at the University of California, Davis approved the study. Data from one participant was excluded due to technical complications with the fMRI scanner, one participant was excluded due to a stimulus computer malfunction, two participants were excluded due to poor behavioral performance in the scanner (defined as falling below trained criterion, 85% correct, in the scanner), and one participant was removed from the scanner before the experiment concluded because they did not wish to continue in the study. Prior to data analysis, to ensure data quality, we conducted a univariate analysis to look at motor and visual activation during the task compared to an implicit baseline (unmodeled timepoints when the participant was viewing a fixation cross). Two subjects showed little to no activation in these regions and were excluded from further analysis. The remaining 23 participants (11 male, 12 female, all right handed) are reported here.

### Stimuli and Procedure

Task stimuli consisted of nine common animals, shown in color on a grey background. Subjects were tasked with learning two “zoo contexts”, consisting of animals organized in a specific spatial orientation (Figure 1a). Importantly, animals in both contexts were visually identical, but each context had a distinct spatial organization. Training consisted of three stages per context: 1) map study, 2) exploration, 3) sequence navigation. This was followed by an additional sequence navigation phase that alternated between contexts.

During map study, participants were initially shown an overhead view of one of the zoo contexts (counterbalanced across participants). After studying this picture, participants were asked to reconstruct the location of all the animals by arranging icons on the screen. If participants were not able to perfectly recreate the maze they were shown the picture once more and asked to try again. Next, during the zoo exploration, subjects used arrow keys to move between items in the zoo, starting from the central animal. At the bottom of the screen participants were shown arrows indicating all possible moves from their current location (e.g. Left, Up, Down, Right at the center position of a maze). If participants made an incorrect move (moving outside of the animal maze) they were informed they made a wrong move. Participants were required to visit each of the animals four times before moving on to the next phase. During the sequence navigation phase, participants were shown a cue with a start and goal animal, and had four moves to reach the goal on a given trial. Start and goal animals were always the endpoints of an arm. Participants were trained to 85% criterion before learning the other context. The same training procedure outlined above was repeated for the second zoo context. After learning each of the zoos to criterion, participants completed an additional sequence navigation phase with the same timing as the MRI scanning session.

In the MRI scanner, subjects completed six runs of the sequence navigation task (Figure 1b). In each run, participants completed 16 sequence navigation trials. Trials from a given context were presented in blocked fashion so that there were 8 consecutive trials from each context. Across runs, context blocks were alternated and their order was counterbalanced across subjects. Each navigation trial began with a cue signaling a start and a goal animal displayed for 3s, followed by a 3s ITI. Subjects then saw the start animal and navigated by pressing buttons to move through the space one animal at a time. Animal items were displayed on the screen for 2s with a 3s ITI, regardless of participant button press. For items where participants made a navigational error, text was displayed for 2s informing them they made a wrong move or incorrectly navigated to a goal animal. In each zoo context, participants planned and navigated 12 distinct sequences (each repeated 4 times across 6 runs of scanning)

### MRI Data acquisition

MRI data were acquired on a 3T Siemens Skyra MRI using a 32-channel head coil. Anatomical images were collected using a T1-weighted magnetization prepared rapid acquisition gradient echo (MP-RAGE) pulse sequence image (FOV = 256 mm; TR = 1800 ms; TE = 2.96 ms; image matrix = 256 x 256; 208 axial slices; voxel size = 1mm isotropic). Functional images were collected with a multi-band gradient echo planar imaging sequence (TR = 1222 ms; TE = 24 ms; flip angle = 67 degrees; matrix=64×64, FOV=192mm; multi-band factor = 2; 3 mm^3^ isotropic spatial resolution).

### MRI data processing

Data were preprocessed using SPM12 (https://www.fil.ion.ucl.ac.uk/spm/) and ART Repair (Mazaika et al., 2009). Slice timing correction was performed as implemented in SPM12. We used the iterative SPM12 functional-image realignment to estimate movement parameters (3 for translation and 3 for rotation). Motion correction was conducted by aligning the first image of each run to the first run of the first session. Then all images within a session were aligned to the first image in a run. No participant exceeded 3mm frame wise displacement. A spike detection algorithm was implemented to identify volumes with fast motion using ART repair (0.5mm threshold) (Power et al., 2012). These spike events were later used as nuisance variables within generalized linear models (GLMs). Subjects native structural images were coregistered to the EPIs after motion correction. The structural images were bias corrected and segmented into gray matter, white matter, and CSF as implemented in SPM12. Native brainmasks were created by combining gray, white matter masks. Data were smoothed with a 4 mm^3^ FWHM 3D gaussian kernel.

### Regions of Interest

ROI definitions were generated using a combination of Freesurfer, and a multistudy group template of the medial temporal lobe. The multistudy group template was used to generate probabilistic maps of hippocampal head, body, and tail as defined by Yushkevich et al. (2015), and warped to MNI space using Diffeomorphic Anatomical Registration Using Exponentiated Lie Algebra (DARTEL) in SPM8. Maps were created by taking the average of 55 manually-segmented ROIs and therefore reflect the likelihood that a given voxel was labeled in a participant. Masks were created by thresholding the maps at 0.5, (i.e., that voxel was labeled in 50% of participants). These maps were then reverse normalized to native subject space using Advanced Normalization Tools (ANTS). Subject specific cortical ROIs were generated using Freesurfer version 6.0. from the Destrieux and Desikan atlas (Desikan et al., 2006, Fischl et al., 2004, Desitrieux et al., 2010). Individual cortical ROIs were binarized and aligned to subjects’ native space by applying the affine transformation parameters obtained during coregistration. These masks were combined into merged masks that encompassed the entire hippocampus bilaterally (see cue period pattern similarity for more information). Anatomical ROIs for V1/V2 and BA4a/p were obtained by running all subjects structural scans through the freesurfer recon-all pipeline. Our V1/V2 ROI was obtained by merging the anatomical masks for BA17 and BA18 (**Fig. S2**).

### Cue period pattern similarity analysis

Our primary interest was to investigate how prospective sequence representations were modulated based on context membership. To achieve this, we used representational similarity analysis to analyze multi-voxel activity patterns (Kriegeskorte et al., 2008) within regions of interest. Generalized Linear Models (GLMs) were used to obtain single trial parameter estimates of the cue period using a modified least-squares all (LSA) model (Mumford et al., 2012, Brown et al., 2016). Data were high-pass filtered using a 128s cutoff and pre-whitened using AR(1) in SPM. All events were convolved with a canonical HRF to be consistent with prior work (Mumford et al., 2012). Cue periods were modeled using separate single trial regressors for each cue (2s boxcar). The remaining portions of the task were modelled as follows: Navigation periods were modelled with separate 25s boxcar functions for each trial, separate single trial regressors for catch sequences modelled as a 15s boxcar, separate single trial catch blank trials (stick function), outcome correct at condition level (stick), outcome incorrect at condition level (stick), and the four button presses at the condition level (stick). Nuisance regressors for motion spikes, 12 motion regressors (6 for realignment and 6 for the derivatives of each of the realignment parameters) and a drift term were included in the GLM. Pattern similarity between the resulting beta images were calculated using Pearson’s correlation coefficient between all pairs of trials in the experiment. Only between run trial pairs were included in the analysis to avoid spurious correlations driven by auto-correlated noise (Mumford et al., 2014).

Based on evidence of functional differentiation along the long-axis of the hippocampus (Poppenk et al., 2013, Bouffard et al., 2021), we tested for any longitudinal or hemispheric differences in hippocampal patterns. Analyses revealed no significant differences in the pattern of results between left and right or between anterior or posterior segments of the hippocampus. As a result, subsequent analyses were performed with pattern similarity data from a bilateral hippocampus mask.

### Linear mixed models

Behavioral responses and pattern similarity were analyzed using linear mixed effects models to account for the nested structure of the dataset, allowing us to statistically model errors in our model clustered around individuals and trial types that violate the assumptions of standard multiple regression models. Statistical comparisons were conducted in R (3.6.0) (https://www.r-project.org/) using lme4 (Bates et al., 2015) and AFEX (Singman et al., 2016). Reaction times were analyzed using the following formula:

EQ1 (Figure 1): RT ∼ Position + (1|subject)

Where (1|x) indicates the random intercept for subject and RT is the reaction time for each position during the navigation phase, excluding position 5 (as no response is required). Furthermore, outlier RTs were excluded that exceeded 2.5 standard deviations from a participants average reaction time.

For the pattern similarity analyses, pairwise PS values were input for each subject into three separate models with the following formulas:

EQ2 (Figure 2b): PS ∼ same_sequence*same_context + (1|subject)

EQ3 (Figure 2c/d): PS ∼ overlap*same_context + (1|subject)

EQ4 (Figure S2): PS ∼ move*same_context + (1|subject)

Where (1|x) indicates the random intercept for subject and PS is the Pearson correlation coefficient for a given trial pair. Fixed effects for EQ1: (1) same sequence - a categorical variable with two levels indicating if the trial pair was from the same or different sequence. (2) Same context - categorical variable with two levels: same or different. Fixed effects for EQ3: overlap - a categorical variable with four levels: full, converging, diverging, and no overlap. Same context - same as EQ2. Fixed effects for EQ4: Move - a categorical variable with three levels: same moves, shared moves, no moves. Same context - same as EQ2. Statistical significance for fixed effects was calculated by using likelihood ratio tests, a non-parametric statistical test where a full model is compared to a null model with the effect of interest removed. For example, to test the significance of an interaction term two models would be fit. One with two main effect and no interaction and the other with the interaction term. Follow up tests and estimated marginal means (Searle et al., 1980) from LMMs were calculated using the R package emmeans (https://cran.rproject.org/web/packages/emmeans/index.html).

In all the above models, a model with a maximal random effects structure, as recommended by Barr et al., 2014, was first fit. In all cases the maximal model failed to converge or was singular indicating over-fitting of the data. When examining the random effects structure for these models, random slopes for our fixed effects accounted for very little variance when compared to our random intercept for subject. To improve our sensitivity and avoid over-fitting these terms were removed as suggested by Matuschek et al., 2017. Lastly, it is important to note that our results are not dependent on using linear mixed models. Using a standard repeated measures ANOVA produces qualitatively and quantitatively similar results in all ROIs.

### Successor Representation Simulation

To better understand specific predictions of the successor representation in our task (Stachenfeld et al., 2017) we performed a simple simulation with respect to our task. First, we created a topological structure (connected graph) that was similar to our task. As seen in **Figure S1**, this structure closely resembled the plus maze participants navigated in. We simulated the successor representation based on a random walk policy using the equation.

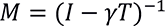

Where *γ* is a free parameter that controls the decay of the SR and T is the full transition matrix of the task depicted in **Fig. S1A/B**. For the current simulations, gamma of 0.3 was used, but results are qualitatively similar for different values. Random walk or policy independence can be assumed in this case because maps were well learned before the scanner and each sequence was traversed in both directions an equal number of times (Momennejad, 2020).

We then tested the hypothesis that, during planning, the hippocampus encodes the SR of the first position in the sequence (columns of SR). We extracted columns of the SR for three planned sequences ((state 1 -> state 5) (state 6 -> state 5) (state 1 -> 9)) and calculated the similarity (Pearsons) between pairs of trials. The same sequence was calculated by correlating the same sequence with itself. The converging condition was obtained by correlating trials that started at different states but converged on the same end state. The diverging condition was obtained by correlating trials that started at the same state but diverged to different end state. Lastly, the no overlap condition was calculated by correlating trial pairs that started and ended at different states. As shown in Figure S1, the SR heavily weights the immediate locations around the starting location and thus would predict that diverging sequences should have higher similarity than converging sequences.

### Timepoint-by-timepoint representational similarity analysis

To examine whether participants activated remote timepoints as they navigated through our virtual environments (e.g., activating decision points early in the navigation trial), we used a variant of single trial modeling using finite impulse response (FIR) functions (Turner et al., 2012). This method allowed us to isolate the unique spatiotemporal pattern of activity for a given navigation trial while simultaneously controlling for surrounding time points during the run. We modeled 47 seconds of neural activity with a set of 38 FIR basis functions. Specifically, we obtained a spatial pattern of activity for each of these 38 TRs in our model, which allowed us to compare the similarity of the spatial patterns of activity between timepoints in the navigation phase. Additional regressors were included for motion, however spike regressors were not included in this analysis because they perfectly colinear with an FIR basis sets for each TR. A separate GLM was used for every trial resulting in 72 voxel time series. To examine within trial type similarity (same trial type across repetitions) timepoint-by-timepoint similarity matrices were generated by correlating activity patterns from repetitions of specific sequence pairs (e.g. zebra-tiger repetition 1 with camel-tiger repetition 1), at every TR. The resultant matrices were symmetrized by averaging across the diagonal of the matrix using the following equation: (X^T^ + X)/2. The resultant timepoint-by-timepoint similarity matrix was averaged within a specific trial type to get a single average timepoint by timepoint similarity matrix for each subject and condition (Fig. 3). This was done separately for converging and diverging sequences. Only between run trial pairs were included in the analysis to avoid spurious correlations driven by auto-correlated noise (Mumford et al., 2014). This method allowed us to isolate individual sequence patterns while controlling for temporally adjacent navigation trials. To identify which points in time corresponded to relevant parts of the task, we manually lagged trial labels by 4 TRs to account for the slow speed of the HRF.

Time point by timepoint similarity matrices were constructed only for converging and diverging sequences. This subset of trials was chosen for several methodological reasons listed below. One is that, to maximally control for differences in trial numbers between conditions and temporally auto-correlated evoked patterns, while still maintaining enough power to examine future state reactivation; we restricted our analyses to converging and diverging sequences within the same context. Importantly, this selection of trials allows us to simultaneously control for several factors while testing specific predictions. Another is that, converging and diverging sequences are matched in terms of the number of shared items and therefore overall visual similarity. Specifically, the same animal items are seen during the first half of diverging sequences, while the same animal items are seen in the second half of converging sequences (all sequences share the center item).

To assess statistical significance, and to correct for multiple comparisons, we used cluster-based permutation tests (Marris and Oostenveld, 2007) with 10,000 permutations, with a cluster defining threshold of 0.05 (two-tailed) and a cluster mass threshold of 0.05. Each pixel of a statistical comparison (T-value) was converted into a Z value by normalizing it to the mean and standard error generated from our permutation distributions.

## Author Contributions

Conceptualization, J.C.D., A.C., CR.; Methodology, J.C.D., A.C., D.H., S.A.P., CR.; Investigation, J.C.D., A.C.; Writing – Original Draft, J.C.D., A.C., C.R; Writing – Review & Editing, J.C.D., A.C., S.A.P., D.H., E.D.B., C.R.; Funding Acquisition, C.R.; Supervision, C.R.

## Data Availability

Data and code needed to reproduce findings here are available from the corresponding author upon reasonable request.

## Competing interests

The authors declare no competing financial interests.

## Footnotes

1 Note that if people retrospectively retrieved past states at later positions or prospectively anticipated future states when they were at earlier positions. However, if participants retrieved past states we would expect off-diagonal activity to be higher for diverging sequences (because the first position was common for the diverging sequences).

## Notes

### Competing Interest Statement

The authors have declared no competing interest.

### Summary of Updates

Figures updated, additional citations, and minor tweaks based on editorial comments.

