## Supplemental Figures for "Goal-centered representations in the human hippocampus"

### Supplemental Materials

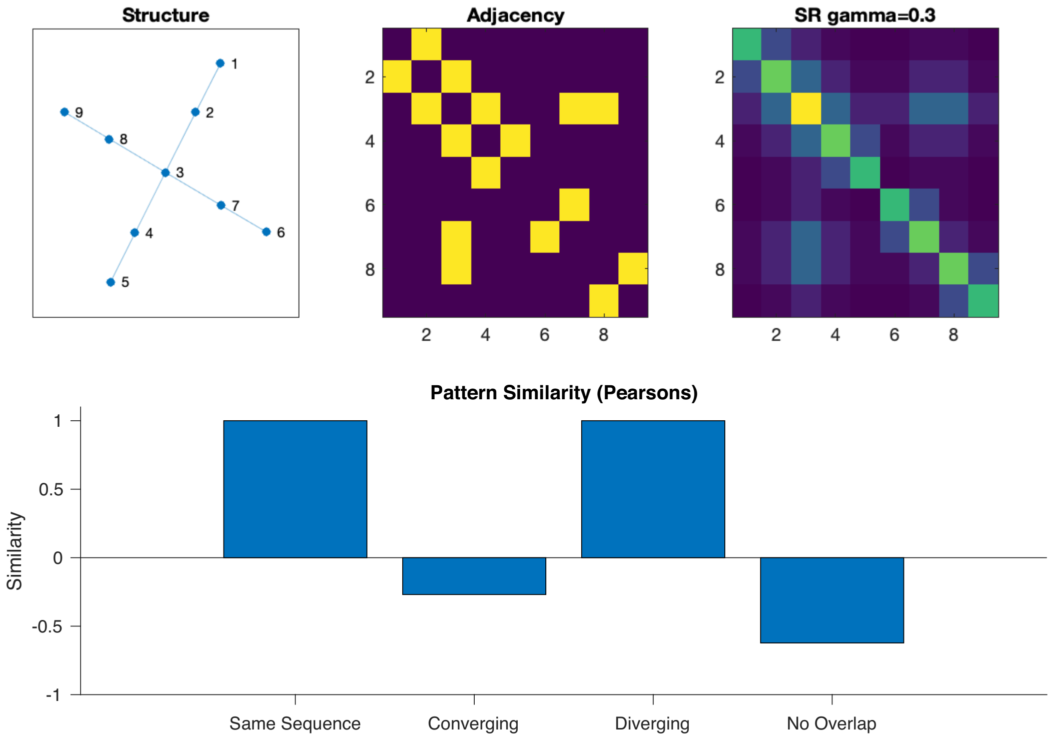


Figure S1: Simulation of Successor Representation (SR) model

Figure S1: Simulation of Successor Representation (SR) model

A) Topological structure of states used in simulation. Numbers identify individual states. Structure is identical to a single zoo context conditions used in our experiment. B) Adjacency matrix of the topological structure. C) SR Matrix using Gamma of 0.3. D) Pattern similarity results. We tested the hypothesis that during planning the hippocampus encodes an SR representation of the first position in the sequence. We simulated three sequences state 1 -> state 5, state 6 -> state 5, and state 1-> state 9. We then indexed the rows of the SR matrix corresponding to the first position in each planned sequence and calculated pairwise similarity using pearsons. Same Sequence = Same Sequence = 1->5 cor 1->5; Converging = 6->5 cor 1->5; Diverging = 1->9 cor 1->5; No overlap 6->5 cor 1->9.

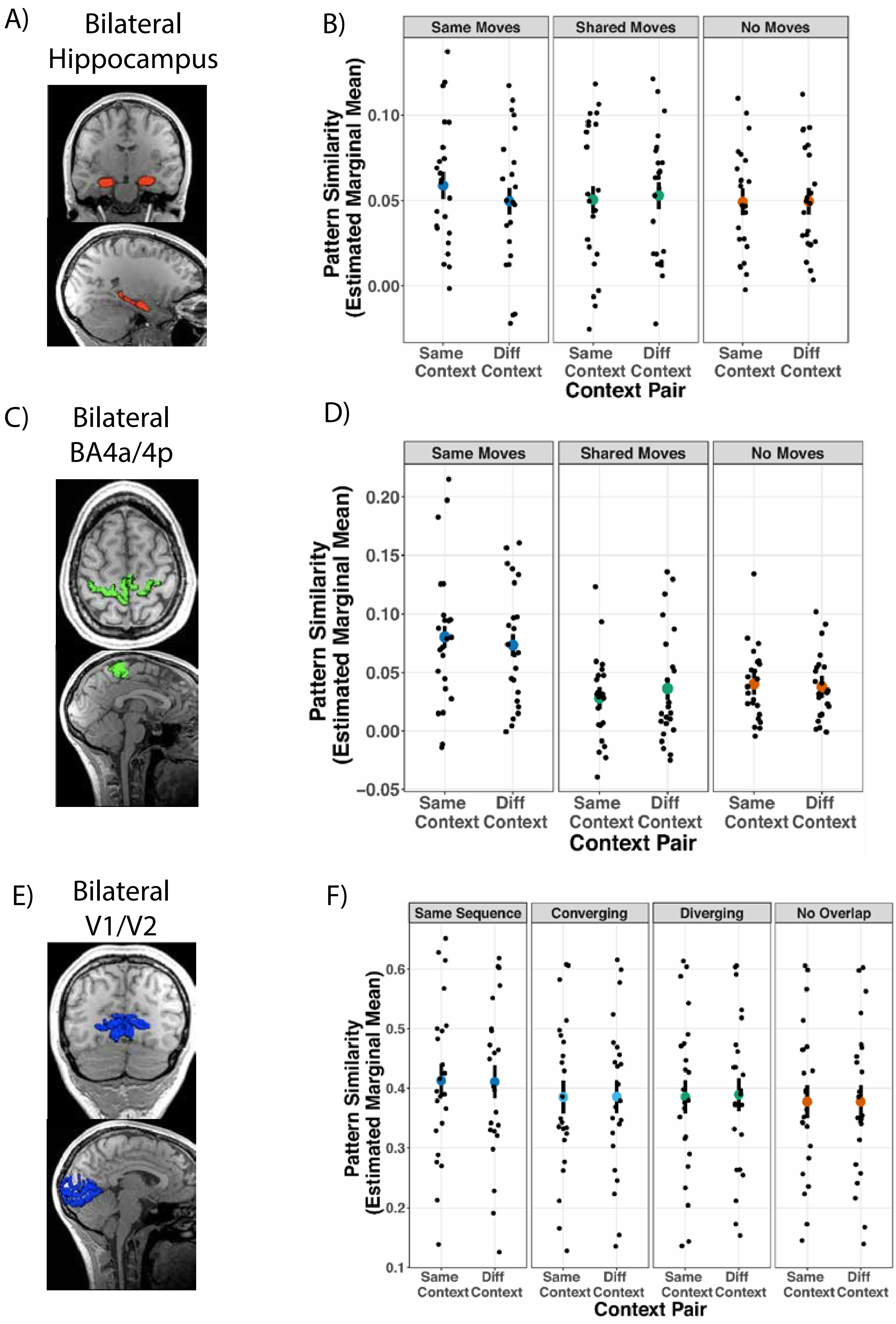


Figure S2: Control analyses in Visual and Motor ROIs

Figure S2: Control analyses in Visual and Motor ROIs

A) Bilateral hippocampal ROI on representative subject's anatomical scan

B) Control analysis contrasting the effect of shared planned motor information on hippocampal representations. Results showed no effect of planned moves or context on pattern similarity (main effect of context: $\chi^2(1, N = 23) = 0.46$, $p = 0.5$; main effect of move: $\chi^2(2, N = 23) = 1.56$, $p = 0.46$; interaction: $\chi^2(2, N = 23) = 2.68$, $p = 0.26$). C) Bilateral primary motor ROI on representative subject's anatomical scan. D) Same as B but examining BA4a/p. $\chi^2(2, N = 23) = 40.40$, $p < 0.0001$, and importantly showed that planned movement was not modulated by context (main effect of move: $\chi^2(2, N = 23) = 40.40$, $p < 0.0001$; main effect of context: $\chi^2(1, N = 23)$, $= 0.01$ $p = 0.94$; Interaction: $\chi^2(2, N = 23)$, $= 1.26$, $p = 0.53$). E) Bilateral primary visual ROI on representative subject's anatomical scan.

F) Control analysis in V1/V2 ROI depicting that visual stimuli is not modulated by context (Main effect of overlap: $\chi^2(3, N = 23\ ) = 90.24$, $p < 0.0001$; main effect of context: $\chi^2(1\ N = 23\ ) = 0.05$; Interaction: $p = 0.82$; $\chi^2(3, N = 23\ ) = 0.76$, $p = 0.86$

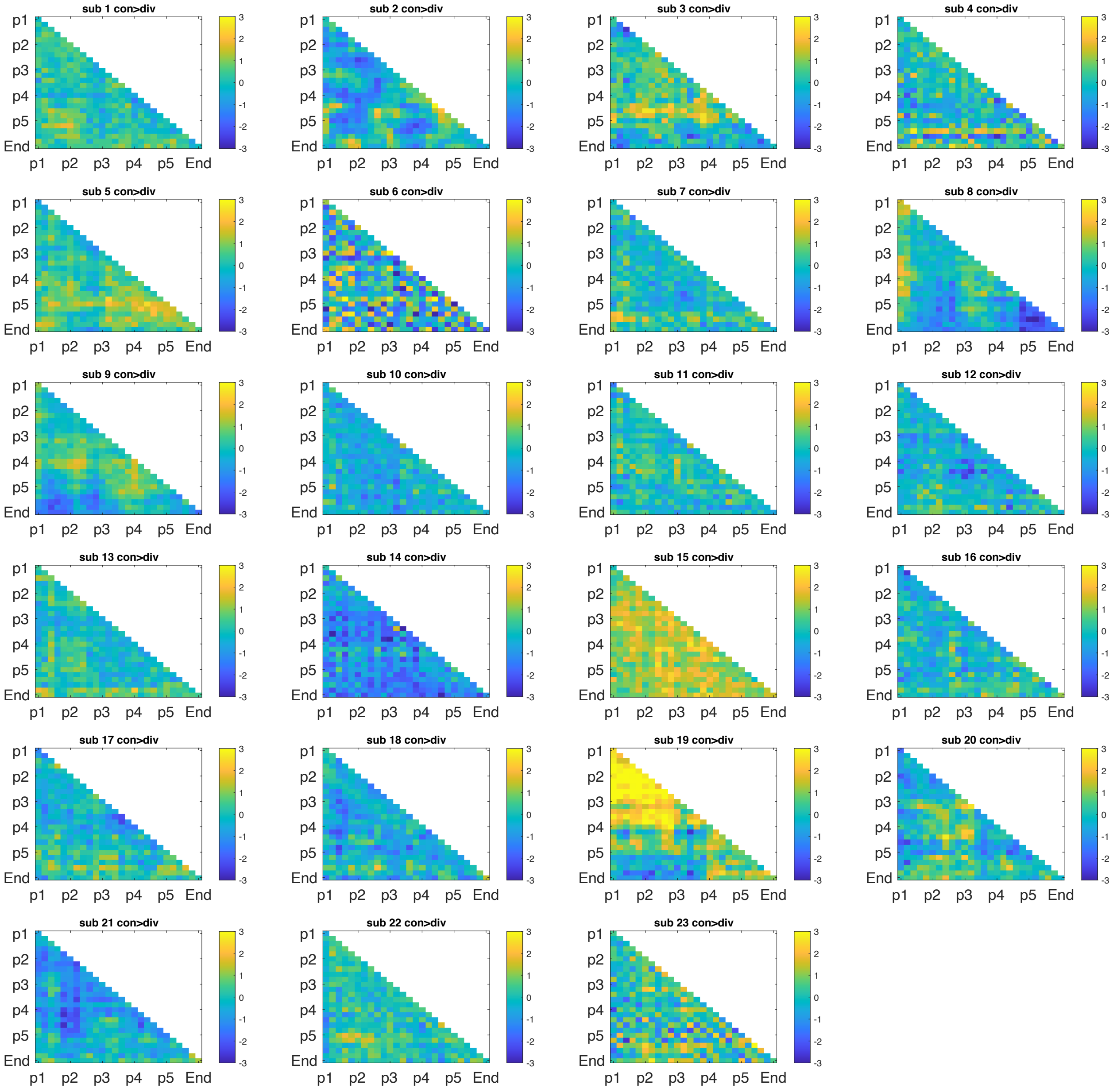


Figure S3: Single subject TR by TR similarity plots for all subjects depicting converging > diverging. Plots are Z scored across subjects all subjects.

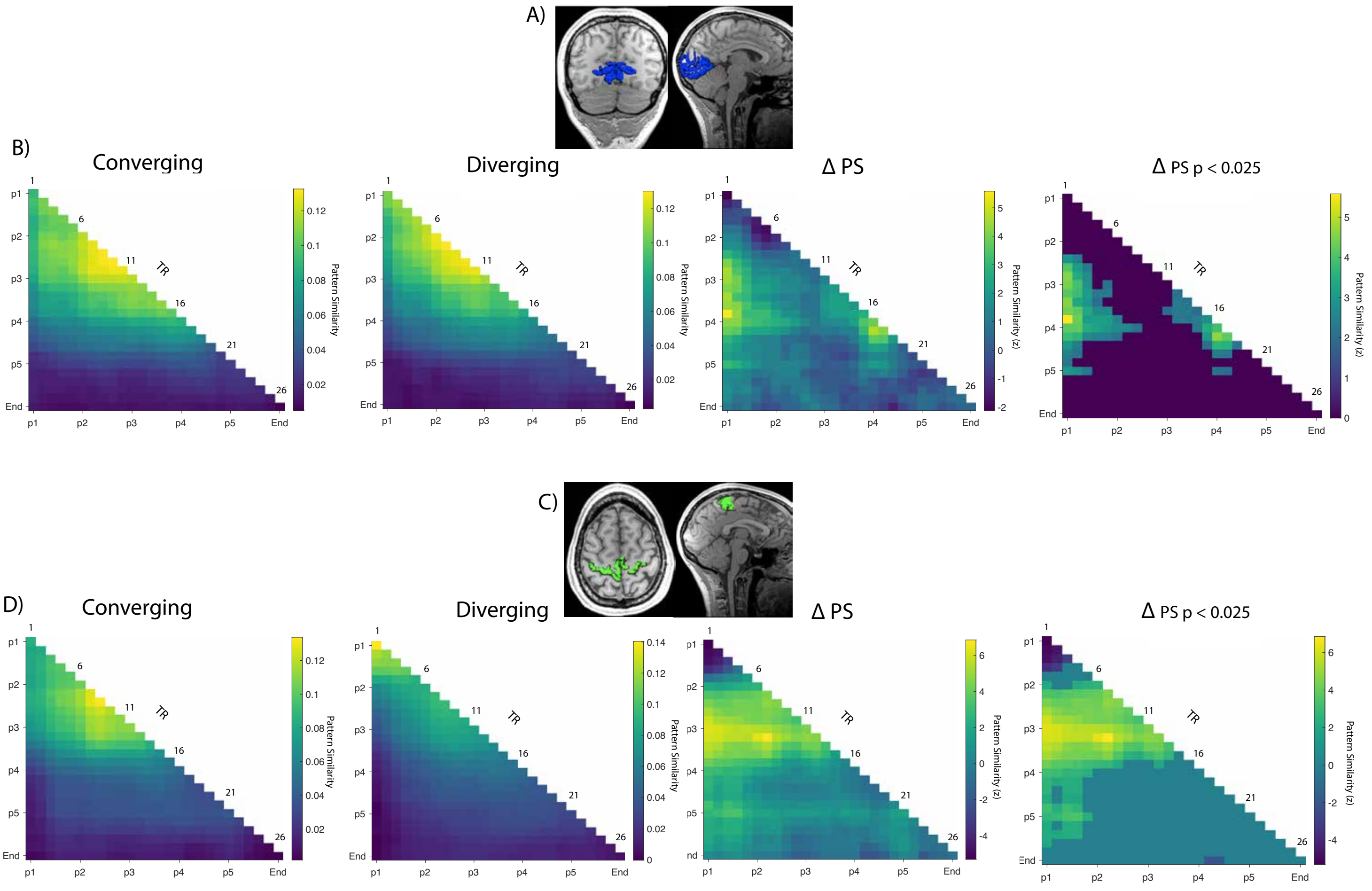


Figure S4 – Visual and Motor regions show off-diagonal reinstatement of upcoming visual and action information

A) Bilateral V1/V2 ROI displayed on representative subject's anatomical scan B) Group level pattern similarity results from converging and diverging sequences during active navigation. ∆PS: TR by TR pattern similarity results depicting a statistical map of converging – diverging. Z values were calculated using a bootstrap shuffling procedure with 10,000 permutations. Thresholded statistical map at $p < 0.025$. Cluster based permutation tests with 10,000 permutations (Maris and Oostenveld, 2007) were performed with a cluster defining threshold of $p < 0.025$ and a cluster alpha of 0.05. C) Bilateral BA4a/p ROI displayed on representative subject's anatomical scan. D) Same as B but in BA4a/p
